# Glby, is a PBP with β-lactamase activity and is required for *in vivo* viability of *M. abscessus*

**DOI:** 10.1101/2021.07.11.451952

**Authors:** Christos Galanis, Emily C. Maggioncalda, Gaurav Kumar, Gyanu Lamichhane

## Abstract

The prevalence of *Mycobacteroides abscessus, Mab*, (also known as *Mycobacterium abscessus*) has been increasing steadily globally. Patients with structural lung conditions such as bronchiectasis, cystic fibrosis and chronic obstructive pulmonary disease are at high risk of developing pulmonary *Mab* disease. The disease is often recurrent as the current treatment regimen is considered sub-efficacious. The cell wall peptidoglycan of bacteria is required for their viability and its biosynthetic pathway is enriched in proteins whose inhibition is the basis for two of the most widely used classes of antibiotics to treat bacterial infections. The peptidoglycan of *Mab* is distinct from that of most bacteria as its synthesis involves penicillin binding proteins (PBP) and L,D-transpeptidases. Here, we demonstrate that *Mab* gene locus *MAB_3167c* encodes a PBP (hereafter referred to as Glby) and is required for normal planktonic growth in liquid broth. Glby exhibits a strong β-lactamase activity and is sensitive to β-lactamase inhibitors. In a mouse model of pulmonary *Mab* disease, mutant lacking this gene was unable to proliferate, gradually cleared and undetectable after three weeks. In a collection of 1.046 *Mab* clinical isolates, there is evidence that changes in amino acid sequence that compromise Glby function are not favored. These evidences suggest that an agent that can inhibit Glby *in vivo* has the potential to be an efficacious treatment against *Mab* disease.

## INTRODUCTION

*Mycobacteroides abscessus* (*Mab*) is a rapidly growing non-tuberculous mycobacteria. It is an opportunistic pathogen that can cause chronic infections with incidences predominantly in patients with structural lung conditions such as bronchiectasis, cystic fibrosis and chronic obstructive pulmonary disease (1, 2). In cystic fibrosis patients, *Mab* is one of the most frequently isolated non-tuberculous mycobacteria (3). Additionally, *Mab* infections of soft tissues in patients that underwent cosmetic surgery have also been reported (4).

*Mab* is considered an emerging pathogen whose incidence in the US exceeds that of tuberculosis and is rising (5). *Mab* infection is largely acquired from the environment and direct person-to-person transmission is rare (6) and considered unlikely (7). A recent report declared *Mab* ‘an environmental bacterium turned clinical nightmare’ (8) citing the following reasons: *Mab* disease is associated with rapid lung function decline and is often incurable (9–11); there are no FDA approved drugs to treat *Mab* disease and cure rate with current treatment based on repurposed antibiotics that need to be taken daily for at least one year is only 30-50% (12); and, *Mab* is naturally resistant to most of the antibiotics available today, and making matters worse the limited selection that can be used to treat this disease is associated with frequent toxicities (13). Current guidelines for treating *Mab* disease are based on clinical experience on repurposing antibiotics, as a regimen informed from a clinical trial has yet to be developed (14, 15). Recent isolates exhibit resistance to an increasing number of antibiotics making treatment of this disease increasingly difficult (16). Despite increasing incidence of this disease, there are critical knowledge gaps in the fundamental biology of this unique organism, which, the emerging evidence suggests, is distinct in many ways from the mycobacteria genus. Even the core genome content is distinct from mycobacteria spp. and was the basis for its recent reclassification as *Mycobacteroides abscessus* from *Mycobacterium abscessus* (17).

Peptidoglycan is the exoskeleton of bacterial cells and is required for their viability, cellular growth and division. A major class of proteins that is involved in the final step of peptidoglycan synthesis in bacteria are penicillin binding proteins (PBP) whose native cellular function is D,D-transpeptidation (18, 19). Additionally, in *M. tuberculosis* for example, the pathway for synthesis of peptidoglycan is enriched in genes essential for its viability, and mimics of metabolites generated in this pathway exhibit antimycobacterial activity (20). The relevance of PBPs in *Mab* to its viability, virulence, and cellular physiology has not yet been described. This study is aimed at fulfilling this critical knowledge gap.

Using bioinformatics approaches we surveyed the genome of *Mab* strain ATCC 19977 (21), which is frequently used as the reference strain in laboratory based investigations. This unbiased survey identified 15 genes with classical PBP features based on sequence homology to known PBPs and the presence of a PBP domain (Table S1). As the biochemical activities and functions of the proteins encoded by these genes have not been characterized, we refer to them as putative PBPs in this study. As it would be beyond the scope of an initial study of *Mab* PBPs such as this one to characterize all putative PBPs, we used known *M. tuberculosis* PBPs as the next criteria to select those that are likely to function as PBPs. *M. tuberculosis* genome encodes for 10 putative PBPs (22–24) which include two protein classes: D,D-transpeptidases and D,D-carboxypeptidases. The D,D-transpeptidases are involved in synthesis of peptidoglycan whereas the D,D-carboxypeptidases catalyze removal of terminal amino acid from peptidoglycan sidechains. The following five PBPs of *M. tuberculosis* belong to the D,D-transpeptidases class: PbpA (Rv0016c), PbpB (Rv2163c), Pbp-lipo (Rv2864c), PonA1 (Rv0050) and PonA2 (Rv3682). These PBPs were used as template to identify homologs in *Mab.* This assessment led to identification of the following genes, *MAB_0035c, MAB_0408c, MAB_2000, MAB_3167c* and *MAB_4901c.* To study these genes, we began by undertaking genetic approaches. Repeated attempts to delete *MAB_0035c and MAB_2000* were made but did not yield any colonies. *MAB_2000* is a homologue of *Rv2163c*, an essential gene in *M. tuberculosis* (25, 26), potentially indicating that it may also be essential in *Mab*. A recent transposon mutagenesis screen was also unable to isolate a *Mab* mutant lacking this gene (27). This screen also reported that loss of *MAB_0035c* affects growth of *Mab*. We were, however, able to generate *Mab* strains lacking *MAB_0408c, MAB_3167c and MAB_4901c.* Growth and colony morphology phenotypes of strains lacking *MAB_0408c* or *MAB_4901c* were unremarkable compared to that of the parent strain. However, strain lacking *MAB_3167c* demonstrated a distinct phenotype in liquid culture, and therefore is the subject of this study. In reference to the ‘globular’ appearance of this mutant in culture and implications of this phenotype to the physiology of *Mab* which is described below, *MAB_3167c* is hereafter referred to as *glby* (Globby). We investigated the relevance of Glby to growth in culture, colony morphology, enzymatic activity against β-lactams, and growth and virulence in a mouse model of pulmonary *Mab* disease.

## RESULTS

### Glby’s amino acid sequence is highly conserved in clinical isolates

*Mab* isolates recovered from patients display significant genomic heterogeneity and differences in antibacterial susceptibility profile indicating that most of the clinical isolates represent a non-clonal collection (28). Availability of a sufficiently large collection of strains would therefore permit assessment of whether specific DNA or amino acid sequences are vital for *Mab* to cause disease in humans. Coding regions that are vital for virulence in humans would be hypothesized to be conserved and any mutation that significantly alters the coded message leading to a loss-of-fitness is likely to be deselected in the clinical collection as it would reduce fitness of the strain in humans. We analyzed the *glby* (*MAB_3167c*) locus in the genomes of 1,046 independent clinical isolates from across the world that are archived in the publicly accessible database PATRIC (29). This database included origins of the strains, which permitted analysis of genomic variations in different regions of the world (Figure S1). Of the 110 SNP locations, only 18 resulted in an amino acid substitution (83.6% were silent mutations) and only one of those (1525A>**G**) was present in >5% of 1,046 strains (Figure 1). This substitution results in a conservative missense mutation from an isoleucine to a valine, **I**509**V**, which was present in 240 strains (22%) and was not endemic to a particular geographic region (**Table 1)**, suggesting independent evolution likely arising from selective pressure. For the countries that had enough representation, only Australia had a low percentage (6%) of strains that harbor this mutation as compared to the UK (22%), the US (25%) and China (25%). The remaining countries did not have an adequate number of strains to enable a statistically accurate representation of strains that harbor this mutation. The second most prevalent amino acid substitution (49 strains or 4.6% of 1,046 strains) is also a conservative missense mutation, A78V, resulting from 233C>**T**. Since the most prevalent amino acid substitutions (I509V & A78V) are conservative substitutions that are considered to produce little or no change in protein function, these results demonstrate that there is evidence of selective pressure *against* changes in the amino acid sequence that may compromise Glby function.

**Figure 1.**
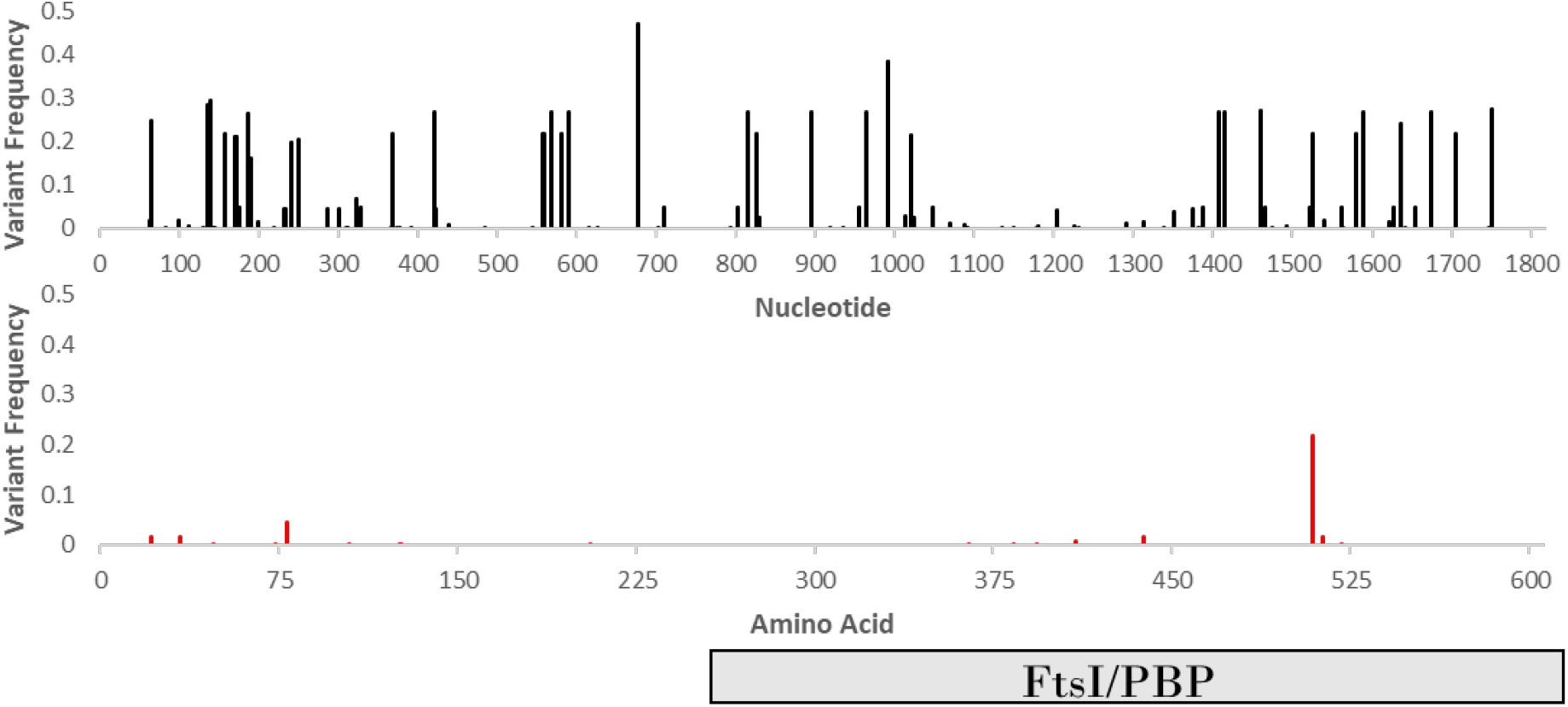
Frequency of mutations in *glby* in clinical isolates. Top-panel, frequencies of single nucleotide polymorphisms (SNP) and, bottom-panel, frequencies of amino acid substitutions in Glby. Each bar represents mutation at the specified location. Predicted FtsI/PBP represents the domain characteristic of PBPs.

**Table 1.** Distribution of I509V mutation in Glby in *Mab* clinical isolates

| Country | # of Isolates | # of isolates with I509V mutation (%) |
| --- | --- | --- |
| United Kingdom | 495 | 107 (22%) |
| China | 275 | 68 (25%) |
| United States | 165 | 42 (25%) |
| Australia | 72 | 4 (6%) |
| Malaysia | 12 | 8 (67%) |
| Denmark | 9 | 1 (11%) |
| Brazil | 6 | 5 (83%) |
| France | 6 | 2 (33%) |
| South Korea | 3 | 2 (67%) |
| Ireland | 3 | 1 (33%) |
| Total | 1,046 | 240 (22%) |

### *Mab* lacking *glby* exhibits altered growth *in vitro*

We used *Mab* ATCC 19977 (hereafter referred to as wild-type, WT), as the parent strain as it is commonly used as a laboratory reference strain (21). Using this strain, we generated a strain lacking *glby* (hereafter referred to as *Δglby*) using a recombineering system optimized for mycobacteria (30). The genome of *Δglby* was sequenced and compared to parent WT genome to verify deletion of the targeted region in *glby* and lack of mutations elsewhere in the genome (Figure S2). We generated a complemented strain by inserting a copy of *glby* cloned from WT into the *attB* site of the *Δglby* chromosome (*attB::pMH94apra-MAB_3167c*) (Figure S3), using pMH94 (31) with modifications to carry an apramycin selection cassette (Figure S4). The genotype of this strain and the site of *glbY* integration was also verified by sequencing its genome (Figure S5). This strain is hereafter referred to as COMP. When grown on Middlebrook 7H10 agar base, colonies of *Δglby* appeared ~ 2 days following the appearance of WT and COMP colonies. The colony morphotype, as assessed by gross visualization of the surface architecture, namely color and size of colonies on 7H10 agar, was not distinct between *Δglby* and that of the WT and COMP strains (Figure S6). However, when grown in Middlebrook 7H9 liquid broth, *Δglby* exhibited a distinct growth morphotype. *Δglby* grew as a single globular clump at the bottom of the culture vessel while the broth was clear, whereas the WT and COMP grew planktonically and turned the broth turbid (Figure 2A). Owing to the ‘globular’ appearance of this strain in liquid broth, we assigned the gene locus (*MAB_3167c*) a phenotype-based annotation *glby* (globby). The reversion of globby growth phenotype to planktonic growth in the COMP strain demonstrates that this phenotype resulted from the lack of Glby. Interestingly, growth of *Δglby* is only permissible in a globular form as ascertained from two distinct experiments. First, we subjected *Δglby* to incremental sheer forces when growing in liquid broth by altering the diameter of the culture vessel while maintaining orbital shaking speed constant at 200 rounds per minute (RPM). When grown in a tube with 1.5 cm diameter (14 mL culture tube, Falcon), *Δglby* grew as a large single globular clump. In a tube with 3 cm diameter (50 mL culture tube, Falcon), *Δglby* grew in several (~ 20-30) smaller clumps and in a 150 mL flask with a 6 cm diameter, *Δglby* failed to grow for five days but formed one large clump after 3 days when the shaking speed was lowered to 75 RPM. In the second experiment, we tested the hypothesis that *Δglby* requires clumped morphotype to sustain growth in liquid broth. We dispersed a *Δglby* culture with vigorous shaking and pipetting into a planktonic suspension and incubated it at standard growth conditions of 37 °C and 200 RPM orbital shaking. *Δglby* again formed globular clumps within 24 hours of dispersal and the broth was clear demonstrating that planktonic growth is impermissible in the absence of *glby.*

**Figure 2.**
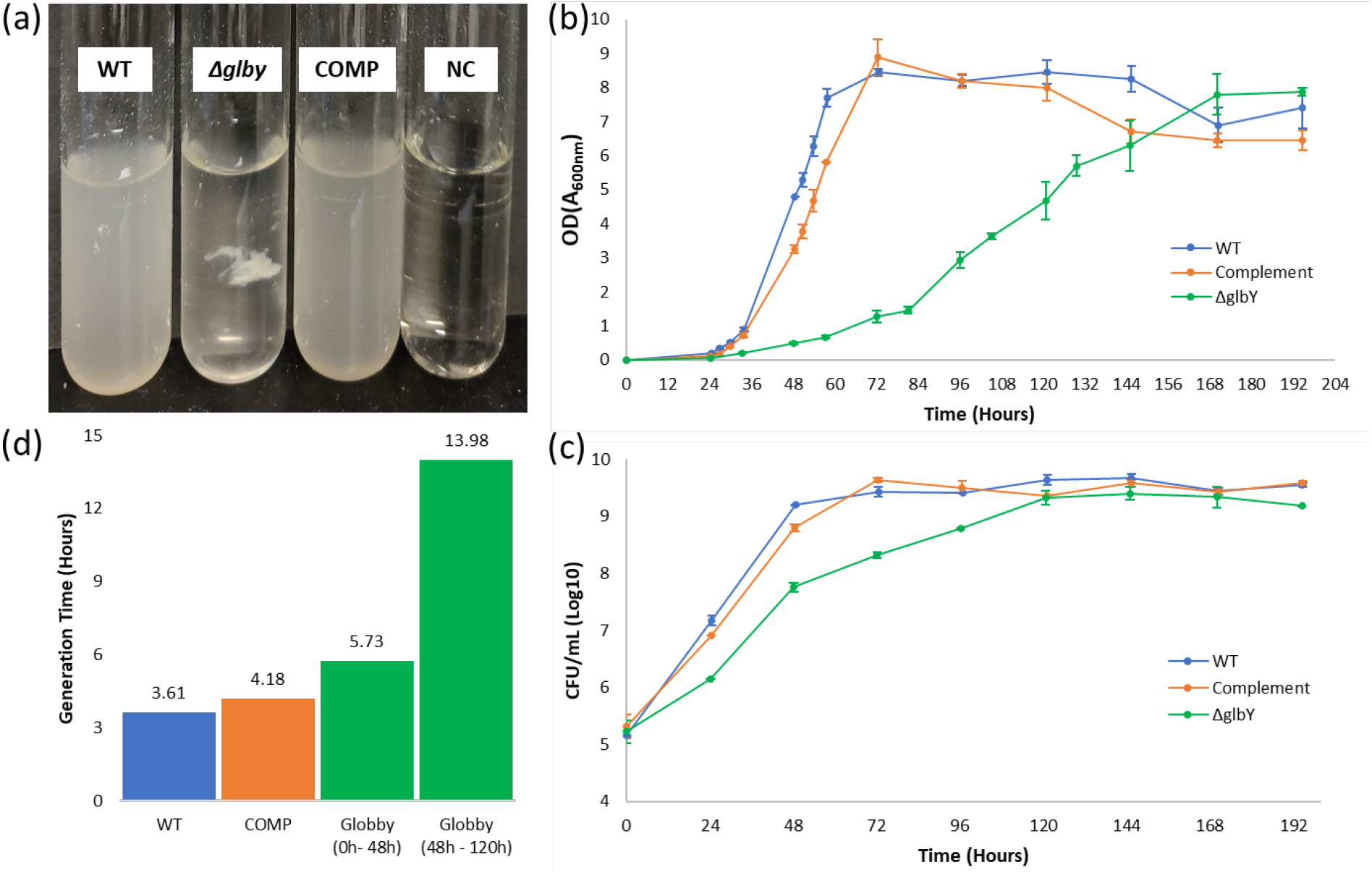
*In vitro* growth phenotypes of *Δglby*. (a) Growth of parent strain *Mab* ATCC 19977 (WT), *Δglby* and Complement (COMP) strains in Middlebrook 7H9 broth. Culture media only as negative controls (NC) was also included. (b) Time course of optical densities of cultures of WT, *Δglby* and COMP measured as absorbance at 600nm. (c) Time course of colony forming units (CFU) of the three strains. (d) Generation time (time required for a strain to double in CFU) of each strain determined from exponential phase(s) of growth. Two generation times reported for *Δglby’s* represent the biphasic exponential growth stages of this strain.

Based on the altered growth phenotype in liquid culture, we hypothesized that the generation time (the time it takes for colony forming units (CFU) to double) of *Δglby* may be different compared to that of WT. To test this hypothesis, *Δglby*, WT and COMP strains were grown under identical conditions and culture optical density (OD) and CFU were determined at regular intervals over 8 days duration (Figures 2B and 2C). Importantly, because of *Δglby*’s globular colony morphology, OD readings would be inaccurate if not dispersed into planktonic suspension. In addition, the OD determinations would likewise be inaccurate if culture from the same tube was sampled at each successive time-point as these interventions may disturb the natural course of *Δglby* growth. Therefore, four culture tubes per time point per strain were included and each sample tube was used only once for OD and CFU determination. *Δglby* exhibited attenuated growth compared to WT and COMP in both OD and CFU determinations (Figure 2B and 2C). In the OD measurement assay, *Δglby* required 168 hours to attain peak OD whereas WT and COMP strains reached peak OD within 72 hours. Interestingly, when *Δglby* reaches at OD of ~4-5 at about 120 hours of growth, which also coincides with the time-point at which it reaches peak CFU density, the broth began turning turbid as it appeared *Δglby* was sloughing off the clumps. At this time, *Δglby* existed in both clumped and suspension forms.

When the growth profile of *Δglby* was compared with WT and COMP in terms of CFU, clear distinction emerged from the onset of growth until peak with largest difference at the 48 hour time-point at which CFU density of *Δglby* was ~1.5 Log_10_ lower compared to WT and COMP strains (Figure 2C). While WT and COMP attained peak CFU density by 72 hours, *Δglby* required an additional 48 hrs (120 hr time-point) to reach the same peak CFU. Of note, *Δglby* appeared to exhibit bi-phasic rates during the exponential stage (Figure 2C), with higher rate of growth between 0-48 hours and reduced rate thereafter until peaking at 120 hours (Figure 2D). In summary, these findings demonstrate that *glby* is required for growth in planktonic form and for normal growth rate; however, it is dispensable for viability or growth of *Mab.*

### Glby is a PBP with strong β-lactamase activity and is inhibited by avibactam

An unbiased query based on pair-wise sequence alignment using BLAST (32) to identify a homolog of Glby identified a protein in *Mycobacterium tuberculosis* encoded by gene locus *Rv2864c*. Glby and Rv2864c share 64% amino acid sequence identity, 78% sequence alignment positivity and an e-value of 0. The *C-*terminus of Rv2864c comprises of a predicted FtsI domain based on which this protein is annotated as a putative PBP. As Glby also possesses the same FtsI domain we hypothesized that Glby may also possess penicillin binding activity. To test this hypothesis, we overexpressed and purified Glby to homogeneity and performed validated assays to verify its ability to bind to penicillins. One of these assays is based on binding to a fluorescent penicillin substrate, Bocillin-FL, which is commonly used to identify proteins with PBP activity (33). We were unable to detect Glby-Bocillin-FL complex while a known PBP of *M. tuberculosis*, PbpB, included in this assay as control exhibited Bocillin-FL binding activity as expected (Figure 3A). There are two possibilities for this observation: 1) Bocillin-FL is not a substrate for Glby and therefore it does not bind to it, which signifies that Glby may not be a PBP, or 2) Bocillin-FL is a substrate of Glby, is bound, but is hydrolyzed rapidly and therefore undetectable as a complex. In order to differentiate between the two possibilities, we used a chromogenic probe, Nitrocefin, that is used to determine if a protein hydrolyzes β-lactam ring such as that present in Bocillin-FL (34). Glby hydrolyzed nitrocefin at a rate comparable to BlaC, the most active β-lactamase in *M. tuberculosis* (35) (Figure 3B, Figure S7). To determine if Glby could hydrolyze a wide spectrum of β-lactams, we incubated Glby with various penicillins, cephalosporins and carbapenems and spectroscopically monitored opening of the β-lactam ring. The penicillins were hydrolyzed most effectively by Glby, followed by cephalosporins whereas carbapenems were least hydrolyzed (Figure 3C). To further probe the ability of Glby to bind β-lactams, we measured Glby’s nitrocefin hydrolysis activity following preincubation with β-lactams (Figure 3D). If Glby is irreversibly bound by a β-lactam, it will exhibit reduced nitrocefin hydrolysis rate. In comparison to the control reaction in which Glby was incubated with nitrocefin only, meropenem, doripenem and biapenem produced the largest reduction in nitrocefin hydrolysis, and several other β-lactams from penicillin, cephalosporin and carbapenem subclasses also affected Glby’s nitrocefin hydrolysis activity as well. Next, we assessed if Glby’s β-lactamase activity is susceptible to inhibition by β-lactamase inhibitors. For this assay we monitored nitrocefin hydrolysis in the presence of β-lactamase inhibitors clavulanic acid, sulbactam, tazobactam and avibactam. These agents are frequently used in combination with β-lactams to treat infections with bacteria that produce β-lactamases. While clavulanic acid, sulbactam and tazobactam reduced β-lactamase activity of Glby by ~50%, avibactam inhibited Glby’s activity by ~91% (Figure 3E). Based on these evidences we conclude that Glby is a PBP, has strong β-lactamase activity and is sensitive to β-lactamase inhibitors.

**Figure 3:**
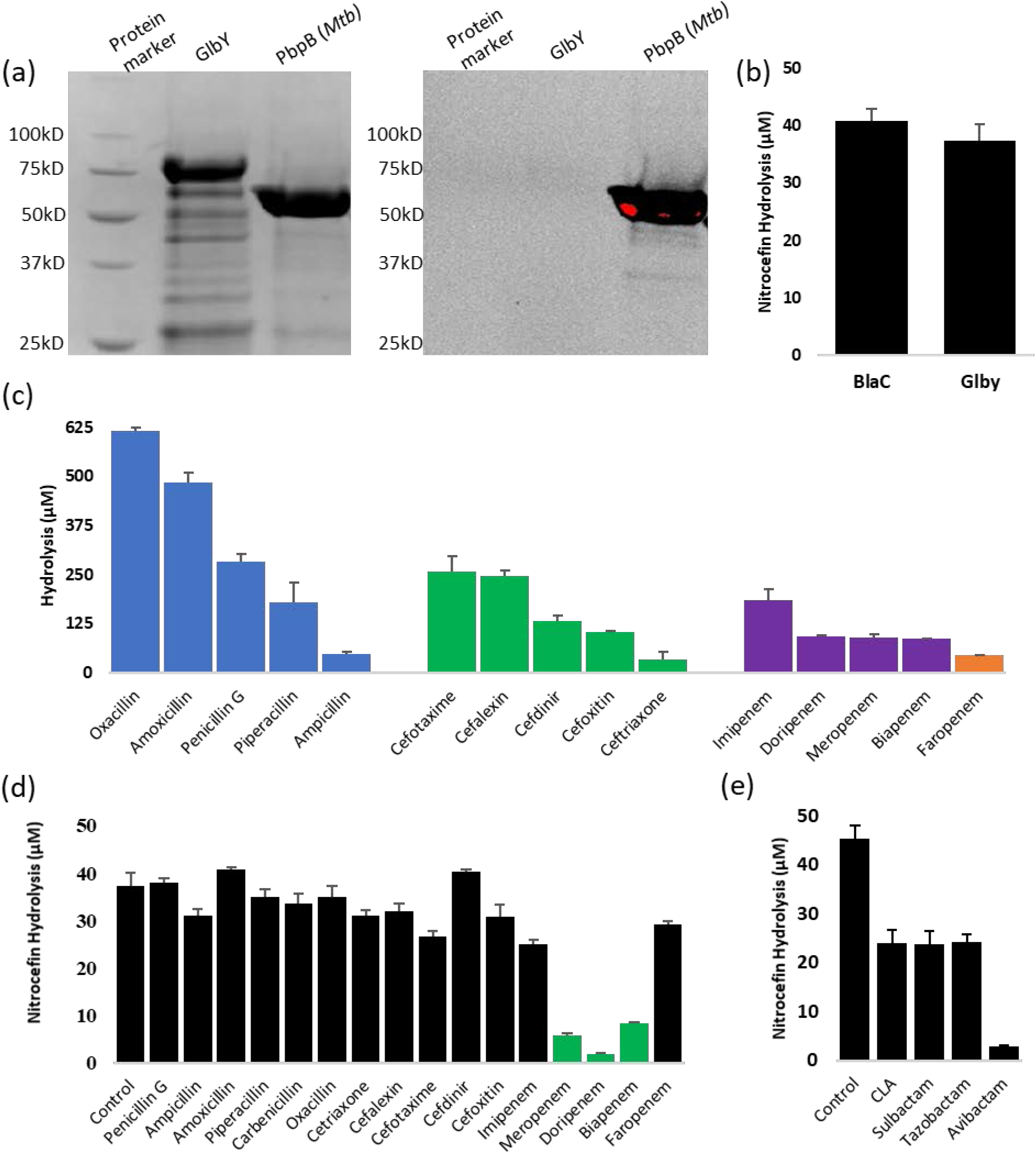
Activity of Glby against β-lactams. (a) Electrophoresis images of Glby and *M. tuberculosis* PbpB (control) incubated with Bocillin-FL. (b) Nitrocefin hydrolysis activities of Glby and a known β-lactamase, BlaC, as control. (c) Hydrolysis of β-lactams by Glby. Inhibition of nitrocefin hydrolysis by β-lactams (d) and β-lactamase inhibitors (e).

### Glby is essential for *in vivo* viability and virulence of *Mab*

On the basis of altered growth rate and morphology, but unaffected viability *in vitro* of *Δglby*, we hypothesized that loss of *glby* would also alter growth and virulence *in vivo* without compromising *Mab* viability. To test this hypothesis, we used a recently developed mouse model of pulmonary *Mab* infection (36) and assessed viability, virulence and growth of *Δglby*, WT and COMP strains. As it is necessary to grow these strains *in vitro* to generate a suspension with which to infect mice, we performed a pilot study to determine the optimal infection dose of *Δglby* required to match the implantation CFU of all three strains due to altered growth and clumping of *Δglby* in culture broth. Although *Δglby* was dispersed prior to infection, it implanted at 10x lower rate than WT and COMP strains (Figure S8). Based on this finding, it was necessary to increase the inoculum of *Δglby* so that following its aerosolization, *Δglby* would implant in the lungs of mice at a burden similar to that of WT and COMP. At 24 hours following infection, the lung burdens of *Δglby*, WT and COMP strains were between 2.8-3.6 Log_10_ and comparable to each other. While the lung burdens of WT and COMP remained steady during the first week, by the third week there was ~2 Log_10_ increase in their CFUs in the lungs of mice (Figure 4). However, lung burden of *Δglby* steadily decreased even from the day of implantation and was undetectable by the three-week time-point although entire lung homogenates from all five mice were inoculated onto growth media for detection of any surviving *Mab.* For instance, by week 2, 99.6% of *Δglby* infection had been cleared as compared to initial implantation levels; by contrast WT grew exponentially to 318x the number of CFUs compared to initial implantation levels. At the four-week time-point, we were again unable to detect any *Δglby* in the lungs of mice, while WT maintained its level and COMP exhibited a slight decrease, but within the standard deviation of WT levels. As a result of not only the *Δglby* CFU burden not increasing, but instead continuously decreasing since the day of infection and reaching undetectable levels, we conclude that Glby is required for viability, growth and proliferation in the lungs of mice. In the study that reported this mouse model, the mice eventually succumbed to death from pathology resulting from the WT strain (36). As mice were able to clear *Δglby* from their lungs, it suggests that Glby is required for virulence of *Mab* to cause lung disease.

**Figure 4.**
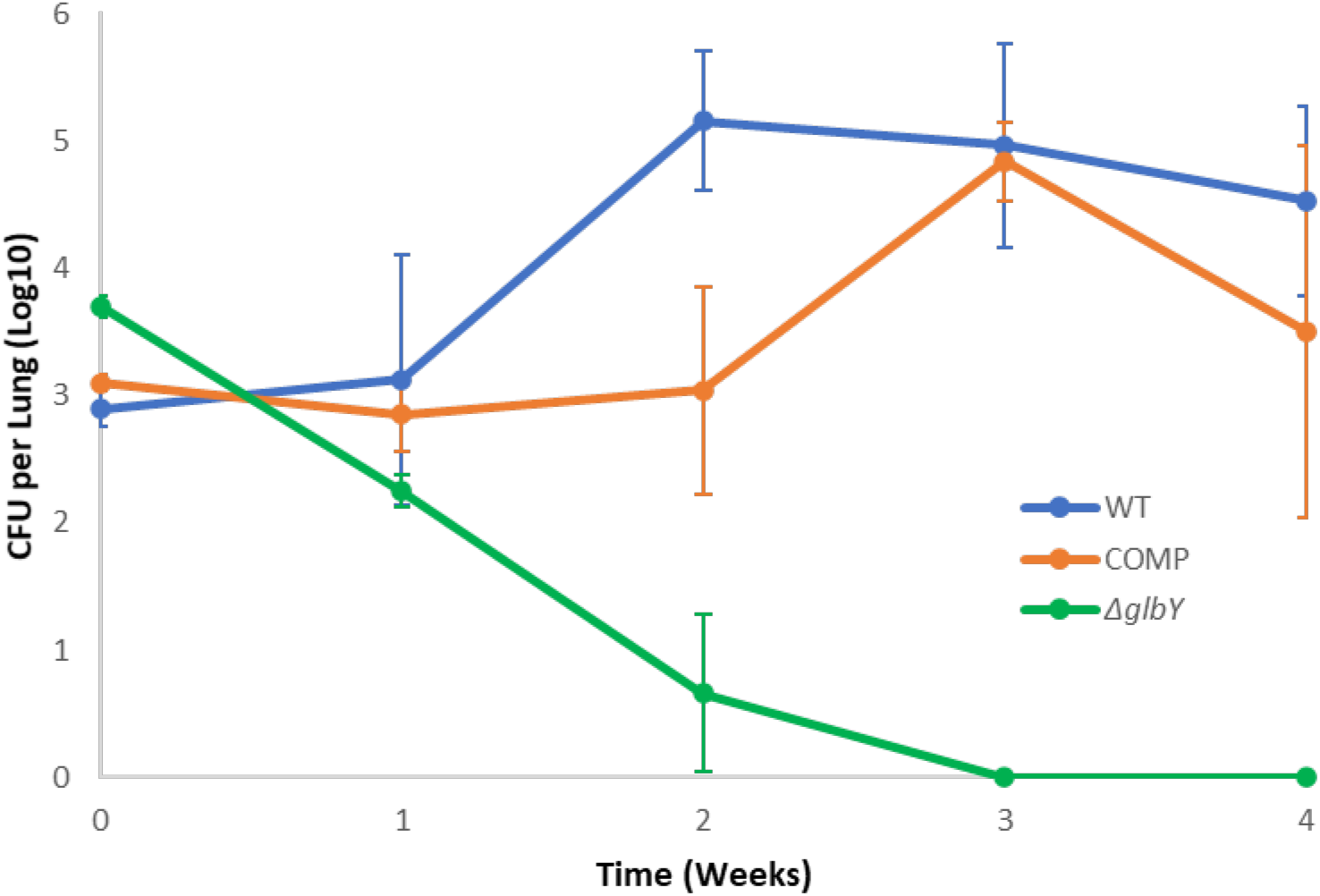
*Mab* burden in the lungs of mice. Burden of *Mab* strain ATCC 19977 (WT), *Δglby* and Complement (COMP) in the lungs of C3HeB/FeJ mice (n=5 per strain per time-point).

## DISCUSSION

The significance of the peptidoglycan biosynthesis pathway to bacteria is underscored by its requirement for viability, growth and division. Considered the Achille’s Heel in bacterial physiology, agents that inhibit peptidoglycan biosynthesis pathway, namely β-lactams and glycopeptides, comprise more than half of all antibiotics prescribed to treat bacterial infections in humans (37). There are significant variations in the chemical compositions and molecular architectures of peptidoglycan among bacteria. Therefore, insights into defining aspects of peptidoglycan can be important in understanding unique physiology of a bacterium.

*Mab* possesses an atypical peptidoglycan characterized by peptide crosslinkages that are distinct from that described in the classical model (18). Unlike in most bacteria, two enzyme classes, L,D-transpeptidases (38, 39) and PBPs with D,D-transpeptidase activity catalyze the final steps of peptidoglycan biosynthesis in *Mab* (40). The findings reported here provide initial descriptions of the relevance of *Mab* PBPs to its viability, growth and disease.

The penicillin binding activity of Glby could be predicted from its sequence homology to the protein encoded by gene locus *Rv2864c*, considered a putative PBP in *M. tuberculosis* (24, 41), and the presence of a FtsI domain that is known to exhibit PBP activity (42). However, its ability to hydrolyze β-lactams from chemically distinct subclasses including penicillins, cephalosporins and carbapenems could not be predicted as only one protein in *Mab*, Bla_Mab_, has been reported to exhibit β-lactamase activity (43). Although β-lactams are active against a wide spectrum of pathogenic bacteria, a majority of β-lactams exhibit poor activity against *M. abscessus* (44). The presence of Bla_Mab_ was considered to be a basis for high level resistance of *Mab* to several β-lactams. Here, we demonstrate that in addition to Bla_Mab_, Glby also exhibits potent β-lactamase activity. The possibility that additional PBPs in *Mab* may also exhibit β-lactamase activity cannot be ruled out. In addition to PBPs, the *Mab* genome also encodes five L,D-transpeptidases (45) that are involved in generating atypical linkages in the peptidoglycan (40). As the L,D-transpeptidases also bind to β-lactams (46) and carry out transpeptidase function, albeit distinct from D,D-transpeptidases (47), it is not known if they also exhibit any β-lactamase activity. A detailed insight into the relevance of all proteins belonging to these two enzyme classes to the metabolism of β-lactams will be necessary to develop a molecular mechanism of the relationship between β-lactams and *Mab.*

PBPs with D,D-transpeptidase activity affect cell wall physiology and consequently the cellular physiology. As bacteria possess several proteins with PBP activity, a direct assessment of each PBP is necessary to identify those relevant to cell wall physiology. The relevance of peptidoglycan synthesis transpeptidases have been described in organisms related to *Mab.* In *M. tuberculosis* and *M. smegmatis*, specific PBPs affect cell morphology (23) and are required for *in vitro* growth (20, 25, 48). Our observations of the aplanktonic, globular *in vitro* growth of *Δglby* is a direct demonstration of its relevance to morphology during growth. In addition, our findings also demonstrate that Glby is dispensable for growth *in vitro* but its loss significantly increases the generation time of *Mab.* A recent study based on generation of a pool of mutants with transposon insertion and detection of outgrowth mutants concluded that *glby* is likely essential for *in vitro* growth (27). In this approach, thousands of mutants were generated together, recovered after growing in a pool and mutants were identified. The genes that are disrupted in the mutants in the pool are predicted to be essential for *in vitro* growth. This approach is a powerful first screen for prediction of essential genes and has high, but not absolute, accuracy of identifying essential genes (25, 26). Essentiality prediction is based on lack of detection of a mutant in the pool, which can also include growth impaired mutants as thousands of other mutants in the pool outgrow impaired mutants and are therefore readily detected in the output. To ascertain if a predicted gene is essential, a direct and site specific mutagenesis is necessary. Therefore, detection of mutations that attenuate *in vitro* growth rate, such as *Δglby*, is a challenge with transposon based random mutagenesis and detection in outgrowth pools. Our findings demonstrate that while Glby is dispensable for *in vitro* growth, it is required for viability and growth in the lungs of mice. Based on this finding, we hypothesize that Glby is required for *Mab* to establish productive infection and cause disease in humans. As the clinical isolates analyzed in our study were obtained from patients, these strains represent *Mab* genotypes that were selected with fitness to cause disease in humans. Therefore, Glby sequences in these mutants represent types that did not compromise viability or virulence of *Mab* in humans. In 1,046 *Mab* clinical isolates analyzed, none of the SNP resulted in nonsense mutations that would lead to a truncated Glby. 83.6% of SNPs were silent and the most frequently observed SNP resulted in conservative amino acid substitution (such as I509V and A78V). As we were unable to identify any deleterious mutations in Glby in clinical isolates, these data strongly suggest that Glby is most likely required for *Mab* viability in humans as well. One approach towards developing a new therapeutic agent is identification of a protein or cellular target in the pathogen that is required for viability or virulence in the host so that, when inhibited, it can arrest the disease and eventually lead to cure it. Using a genetic approach and a mouse model of pulmonary *Mab* infection, we have provided evidence that biochemical inactivation of pathogenic *Mab* with an antibiotic specific to Glby has the potential to exhibit bactericidal activity and improve treatment outcomes.

## MATERIALS AND METHODS

### Bacterial Strains and Growth Conditions

*M. abscessus* strain ATCC 19977 (21) was procured from ATCC (Manassas, Virginia), was used as the parent strain and all strains were derived from it. In general, validated protocols for preparing common reagents for handling and growing mycobacteria were used (49). All strains were grown in Middlebrook 7H9 broth (Difco) supplemented with 10% albumin-dextrose-saline enrichment (ADS), 0.5% glycerol, and 0.05% Tween 80 with constant shaking at 200 rpm in an orbital shaker at 37 °C. To culture on solid media, Middlebrook 7H10 agar (Difco) supplemented with 10% ADS and 0.5% glycerol and appropriate antibiotic (if any), depending on the resistance marker, were used. All inoculations that required agar base were performed on Middlebrook 7H10 with the exception of mouse lung homogenates, which were inoculated onto selective Middlebrook 7H11 agar (Difco) supplemented with 10% ADS, 0.5% glycerol, 20 mg/L Trimethoprim (Sigma-Aldrich, T7883), 50 mg/L Carbenicillin (Fisher Scientific, 50-213-247), and 50 mg/L Cycloheximide (Sigma-Aldrich, C7698). *E. coli* strain DH5α (NEB Labs, C2987H) was used for cloning and *E. coli* strain BL21(DE3) (NEB Labs, C2527) was used for protein overexpression. These strains were grown in LB Broth as specified by the manufacturer.

### Genetic manipulation of *Mab*

For preparation of electrocompetent cells, *Mab* strain ATCC 19977 was grown in Middlebrook 7H9 broth to mid-log phase. A subculture in 100 mL of Middlebrook 7H9 supplemented with 0.2% succinate was initiated and monitored optical density (measured as absorbance at 600nm) until it reached 0.5–0.8 at which point the culture was placed on ice for 30 minutes divided into two 50 mL aliquots. These cells were washed four times as follows: the cell suspension was centrifuged at 3,000 rpm for 10 minutes at 4°C, supernatant was discarded and the pellet was resuspended in 40mL of ice-cold 10% glycerol. After the final wash, the cell pellet was resuspended in 1 mL 10% glycerol and 200 μL aliquots were transferred into microfuge tubes and stored at −80°C until use.

For genetic manipulation of *Mab*, electrocompetent ATCC 19977 was transformed with pJV53 in accordance with the recombineering protocol (30). Briefly, *Mab* competent cells were incubated on ice with ~500 ng of DNA for 10 minutes and transformation was performed with the following settings (2.5kV, 25 uF, 1000ohms) using an electroporator (BioRad). Cells were recovered in 1ml of Middlebrook 7H9 broth incubated for 4 hrs at 37°C and inoculated onto Middlebrook 7H10 agar supplemented with appropriate antibiotics. For selection of pJV53, 128 μg/ml kanamycin was used, for allelic exchange substrate 64 μg/ml Zeocin and for pMH94-Apra, 25 μg/ml Apramycin was used.

### Cloning

All PCRs were carried out using Phusion High Fidelity Polymerase (NEB Labs, M0530). To begin with we PCR amplified *Zeo^R^* cassette flanked by *loxP* sites using pMSG360zeo as template (50) and inserted this fragment into pUC19 at the multiple cloning site. Next, we PCR amplified the ~1,500 bp regions from 5’ and 3’ ends of *glby* using PCR and cloned into the flanks of *Zeo^R^* cassette. This plasmid was used as the template to generate linear allelic exchange substrate spanning 5’*glby*-*loxP-Zeo^R^*-*loxP-3’glby*. pMH94-Apra was generated by replacing Kan^R^ cassette from pMH94 (31) with ApraR gene cloned from pCAP03-acc(3)IV (51) using primers that had a complementing overhang with the PCR product from pMH94 for easy restriction cloning.

For overexpression of Glby, we PCR amplified *glby* using genomic DNA of *Mab* strain ATCC 19977 as template and inserted the amplicon into pET32a vector. The region in the *N-*terminus of Glby that is predicted to encode a transmembrane anchor domain, amino acid residues 1-19, was excluded during cloning to facilitate extraction and solubilization of Glby. DNA sequences of all resulting plasmids were determined via Sanger sequencing (Eurofins, KY) and only clones with desired sequences were selected for study.

### Protein Overexpression and Purification

Glby was overexpressed in *E. coli* BL21(DE3) carrying pET32a-*glby* cells by inducing a 1-liter culture in LB broth with 0.25 mM IPTG overnight at 16°C with orbital shaking at 150 RPM. Glby is expressed from pET32a (52) as a recombinant protein with *N-*terminal fusion of TRX-tag, 6His-Tag for Ni-NTA binding, thrombin and enterokinase sites to facilitate solubility during expression and cleavage of the Glby fragment during purification. Despite the presence of TRX tag, a significant fraction of Glby was present in the pellet. To address this, induced cells were resuspended in sonication buffer (Tris-Cl 50mM, NaCl 300mM, and Imidazole 25mM, pH 8), sarkosyl (1% final concentration) and 1mM PMSF were added and incubated overnight at 37°C with constant shaking at 150 RPM. Cells were then sonicated for 5 mins (intervals of 30 seconds on, 30 seconds off) on ice. Solubilized Glby was purified using the His-Tag via Ni-NTA based metal affinity chromatography using an AKTA FPLC (GE Healthcare, USA). The protein sample was then washed with at least 5 column volumes in washing buffer (identical to sonication buffer except for sarkosyl and PMSF). Glby was eluted with Tris-Cl 50mM, NaCl 300mM, and imidazole 500mM, pH 8 and the 6His- and TRX tags were cleaved with Thrombin. Imidazole was removed by dialyzing using 50mM Tris pH8, 300mM NaCl, 0.1 mM PMSF. From this preparation, Glby was further purified using nickel affinity column to remove thrombin and the TRX- and His-Tags. Glby concentration was determined using a spectrophotometer and 1mg/ml protein aliquots were prepared for assays. *M. tuberculosis* BlaC and PbpB proteins prepared in a previously reported study (53) were used.

### gDNA extraction, Whole Genome Sequencing and assembly

Genomic DNA (gDNA) from all strains was extracted and purified as described for mycobacteria (49). The purity of gDNA preparations were determined spectrophotometrically. Sequences of genomes of WT *Mab* strain ATCC 19977, Δ*glby* and COMP were determined using Illumina PE150 platform (Novogene, CA, USA). We re-sequenced our stock of WT *Mab* ATCC 19977 strain to identify pre-existing SNPs in comparison to the reference sequence of ATCC 19977. The gDNA sequence of our stock of ATCC 19977 was used for comparison with the sequences obtained for Δ*glby* and COMP. To verify genotype of *Δglby*, Geneious v11.1.5 (Biomatters) was used to map the sequence reads obtained from gDNA of this strain to the reference strain (Figure S2). To verify the genotype of the complement strain, *de novo* assembly of the reads was performed, and genes were subsequently annotated in the generated contigs with 100% identity to genes in *Mab* ATCC 19977. Neighboring genes to the *attB* were used to localize the integrated plasmid (Figure S5).

### Distribution of mutations in Glby in *Mab* clinical isolates

Whole genome sequences of 1,046 *Mab* clinical isolates from around the world that are archived in the PATRIC database (29) including their locations of origin were considered in our study. Using Geneious v11.1.5 (Biomatters), the sequence of *glby* from all genomes was extracted by creating a custom BLAST database within Geneious. Using ATCC 19977 *glby* as the reference sequence, all 1,046 genomes were queried and the identified sequences were aligned with MUSCLE alignment tool included in the software and a consensus sequence of *glby* was generated. SNPs and variations in *glby* in each of the 1,046 strains were identified by comparison against the consensus sequence using the ‘Find Variations/SNPs’ algorithm in Geneious.

### Determination of *Mab in vitro* growth profiles

Stocks of *Mab* WT, Δ*glby* and COMP strains, archived in −80 °C, were used to inoculate Middlebrook 7H9 broth to generate primary cultures. These cultures were used to inoculate 120 mL Middlebrook 7H9 broth with the same starting OD of A_600nm_ = 0.0005. This suspension was used to transfer 2 mL aliquots into a 14 mL culture tube. Four distinct tubes/samples were allocated per time point for each strain and OD was determined by measuring absorbance at 600 nm (A_600nm_). Simultaneously, for each strain, 100 μl of appropriate dilutions of each sample at planned time-points were inoculated onto Middlebrook 7H10 medium, incubated at 37 °C for five days for WT and COMP and eight days for Δ*glby* and CFU were enumerated. Δ*glby* CFUs required additional three days to appear compared to WT and COMP strains and thus the difference of incubation duration.

### Bocillin-FL binding and β-lactam and nitrocefin hydrolysis assays

Bocillin-FL (ThermoFisher Scientific, Catalog # B13233), a fluorescent penicillin (λ_max_ = 504 nm) (33), 50 μM final concentration was incubated with Glby and control comparator protein PbpB of *M. tuberculosis*, each at a fixed amount of 10 μg, in Tris-Cl buffer pH 7.4 for 30 minutes in dark at 35°C. The reaction was quenched by adding Laemmli buffer (BioRad, #1610747) and denatured by incubating at 95 °C for 5 minutes. The mixtures were electrophoresed on 14% SDS PAGE and bocillin fluorescence was imaged using Gel Doc EZ imager (BioRad).

To assess β-lactamase activity, Glby (and BlaC of *M. tuberculosis*) 10 μM, was mixed with nitrocefin (Calbiochem) 100 μM, in 10 mM Tris-Cl buffer pH 7.4 in a final reaction volume of 100 μl and incubated at 25 °C for 40 minutes. As a control, nitrocefin in buffer, but without any protein was included. Hydrolysis of nitrocefin was monitored by measuring the absorbance specific to the hydrolyzed product (λ_max_ = 496 nm) (34). To account for the low level of hydrolysis of nitrocefin in the buffer itself, the absorbance in the control reaction at each time-point was used to correct the absorbance readings in reactions containing Glby and BlaC. To assess if nitrocefin hydrolysis activity of Glby is affected in the presence of β-lactams or β-lactamase inhibitors, Glby was incubated with each β-lactam or β-lactamase inhibitor, 1 mM final concentration, at 25 °C for 30 minutes following which nitrocefin, 100 μM, was added. Reaction conditions and monitoring of nitrocefin hydrolysis were undertaken as described above.

Opening of the β-lactam ring in each drug via hydrolysis by Glby was measured spectrophotometrically using the extinction coefficient specific to each β-lactam class (penicillins: 235 nm; cephalosporins: 260nm; carbapenems: 300nm). Penicillin G, Ampicillin, Amoxicillin, Oxacillin, Piperacillin, Cefotaxime, Cefalexin, Cefdinir, Cefoxitin, Ceftriaxone, Iminipenem, Doripenem, Meropenem, Biapenem, Faropenem, Clavulanic Acid, Sulbactam and Tazobactam were purchased from Sigma-Aldrich. Avibactam was purchased from Santa Cruz Biotechnology. All β-lactams were used at a final concentration of 1 mM and mixed with 10 μM Glby in a final reaction volume of 100 μl in Tris-Cl buffer, pH 7.4. Reactions were monitored for 60 minutes at 25 °C using the Spectra Max 250 spectrophotometer (Molecular Devices, USA). Absorbance measurement time course was used to determine the rate of hydrolysis of each β-lactam by Glby.

### *In vivo* viability and growth assessment of *Mab*

A mouse model of pulmonary *Mab* infection and the protocol described (36) was used to assess the requirement of Glby for viability and growth of *Mab.* Briefly, C3HeB/FeJ mice, female, 5-6 weeks old (Jackson Laboratoreis, Bar Harbor, Maine), 25 mice per infecting strain, were infected with an aerosol of cultures of WT, Δ*glby* or COMP obtained at exponential phase of growth and diluted to an OD of A_600nm_ = 0.1 in sterile 1x PBS, pH 7.4 in a Glas-Col nebulizer according to the manufacturer’s instructions (Glas-Col, Terre Haute, Indiana). One week prior to infection and throughout the study, mice were treated daily single dose of dexamethasone, 5 mg/kg/day as specified in the mouse model protocol. Five mice per infection group were sacrificed at one day (week 0), 1, 2, 3 and 4 weeks following infection, lungs were homogenized, inoculated onto Middlebrook 7H11 selective agar, incubated at 37 °C for five days for WT and COMP and eight days for Δ*glby* and CFU were enumerated. Mean CFU ± standard deviation was calculated to determine the CFU burden of each strain over the time course of the study.

## Supporting information

All Supplemental Data

## ACKNOWLEDGMENTS

This study was supported by grants R21 AI137720 and R01 AI155664 to GL. ECM was in part supported by NIH F31 award HL147392.

## AUTHOR CONTRIBUTIONS

GL and CG conceived and devised the study. CG conducted bioinformatics, genetic, cell and microbiology, and animal studies. ECM generated plasmids for genetic manipulation of *M. abscessus.* CG and GK conducted biochemistry studies. GL and CG analyzed data, wrote manuscript and all authors were involved in its revision and finalizing.

