## Supplementary material for "Glby, is a PBP with β-lactamase activity and is required for *in vivo* viability of *M. abscessus*": All Supplemental Data

**SUPPLEMENTARY INFORMATION**

**Table S1. Genes bioinformatically identified with classical PBP Features.**

| | | |
|---|---|---|
| *MAB_0035c* | *MAB_2000* | *MAB_3167c* |
| *MAB_0330* | *MAB_2019* | *MAB_3234* |
| *MAB_0408c* | *MAB_2179* | *MAB_3681* |
| *MAB_0519* | *MAB_2833* | *MAB_4800* |
| *MAB_1870* | *MAB_2875* | *MAB_4901c* |

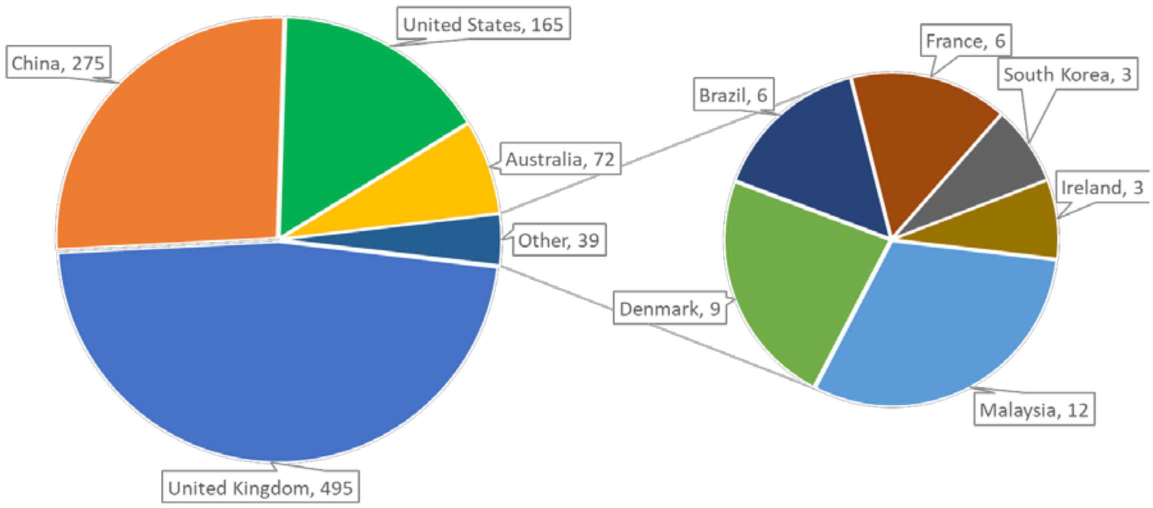


**Figure S1. Geographical origins of *Mab* clinical isolates whose genomes are archived in the PATRIC database.**

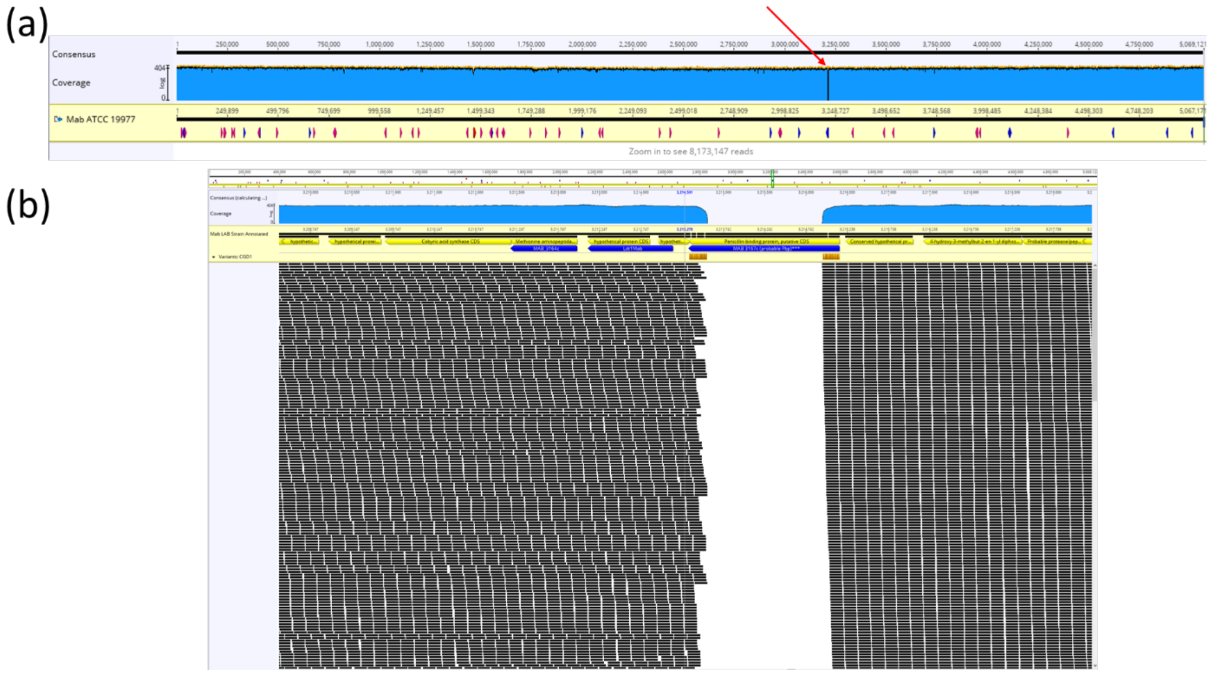


**Figure S2. Mapping and alignment of whole-genome sequencing reads of Δ*glby* to the genome of reference strain.** (a) Coverage of ~8 million WGS reads on *Mab* ATCC 19977 genome. Blue area indicates coverage of reads. (b) Close-up of red arrow location from above (locus MAB_3167c).

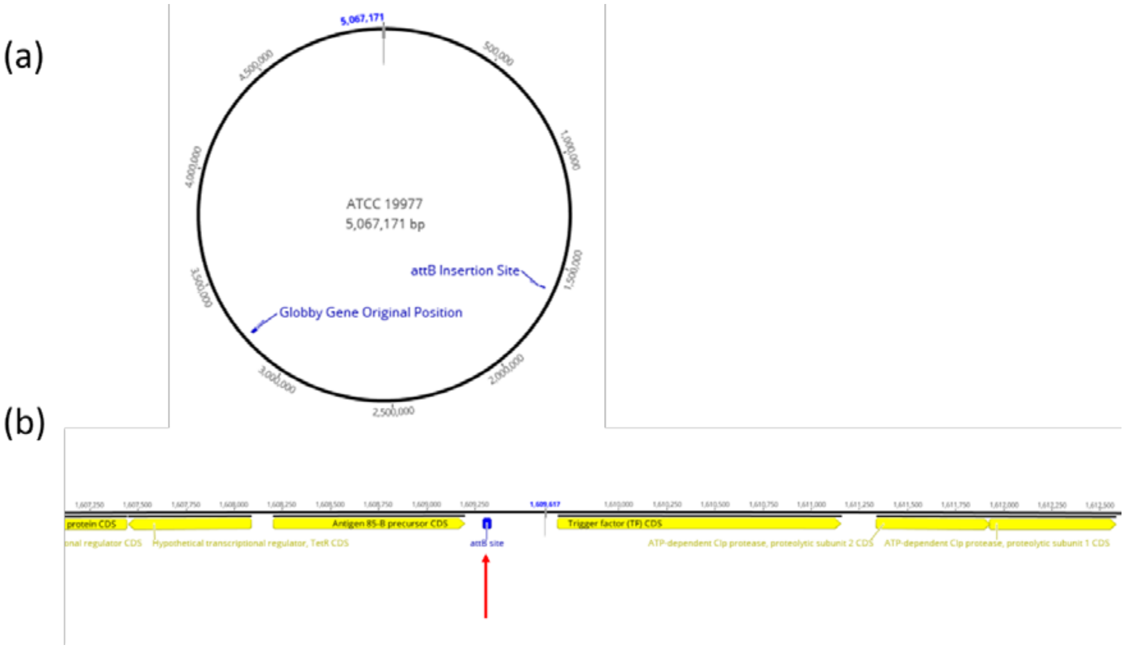


**Figure S3. Genomic Insertion Site for pMH94 based plasmids.** (a) Map of *Mab* ATCC 19977 showing *glby*'s original position within the genome and location of the *attB* insertion site. (b) Close-up of *attB* insertion site. Arrow shows *attB* insertion site in the genome of *Mab* ATCC 19977. (b) pMH94Apra-MAB_3167c plasmid used for complementation.

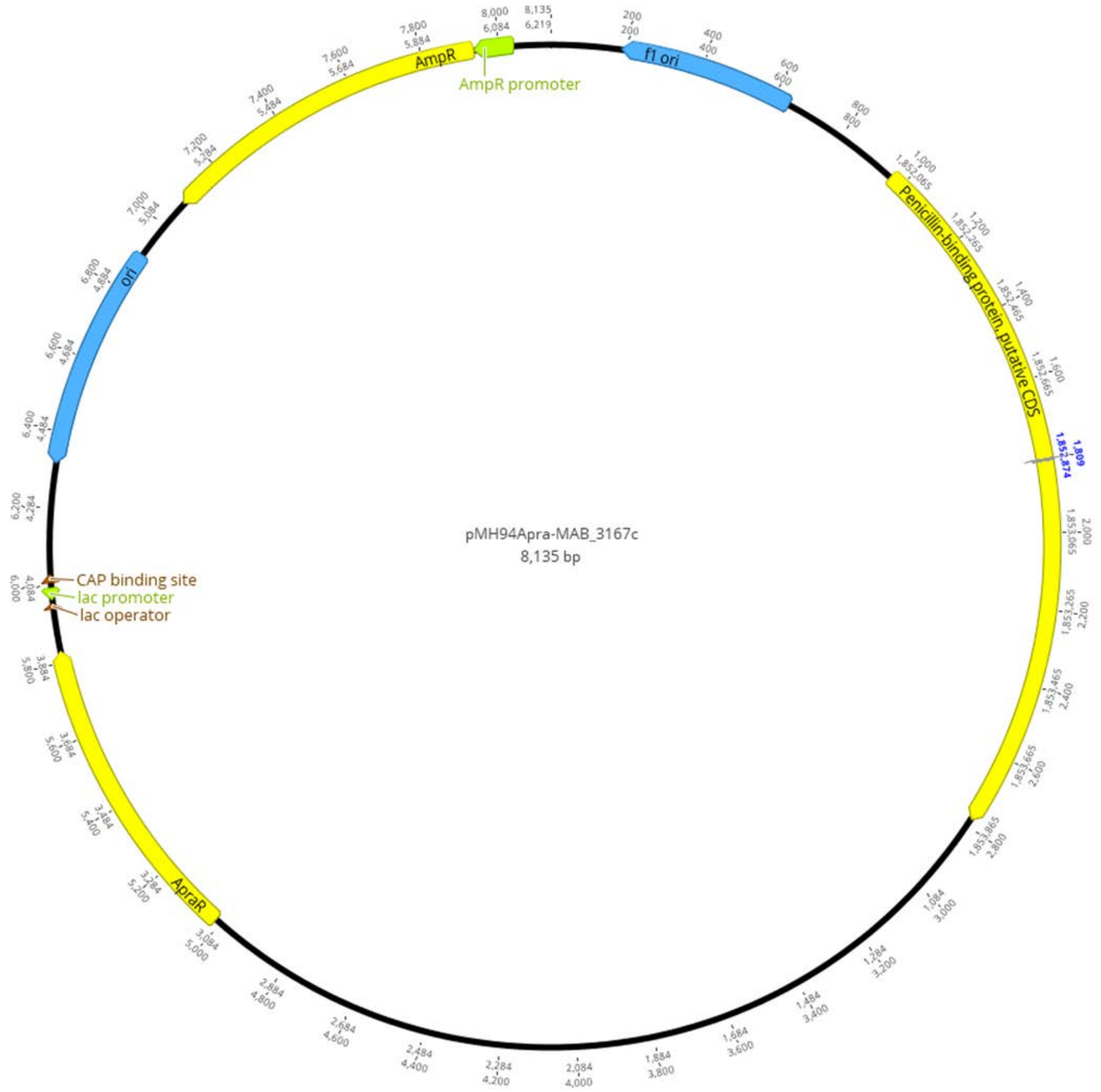


**Figure S4. Map of pMH94Apra-MAB_3167c plasmid used for complementation in *Mab*.**

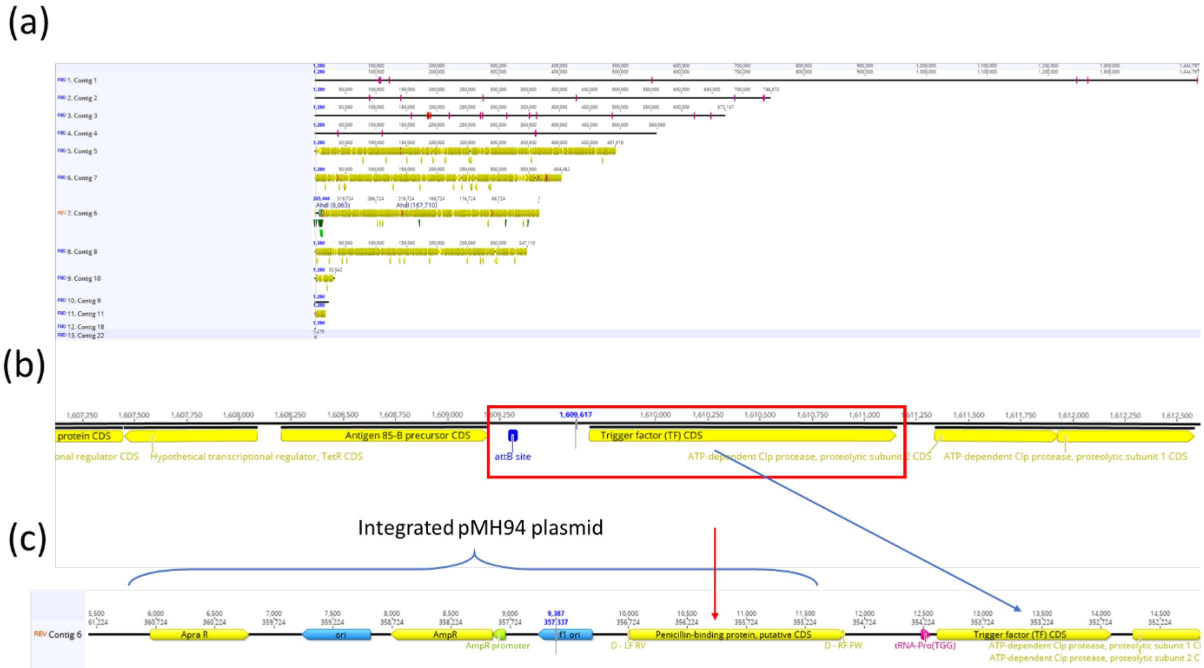


**Figure S5. *De novo* assembly of COMP strain.** (a) *de novo* assembly results of COMP strain which has been fully annotated based on 100% identity to original genes in *Mab* ATCC 19977. (b) Close-up of *attB* insertion site in *Mab* ATCC 19977. Red box indicates location where pMH94 based plasmid is expected to insert and Trigger Factor (TF) CDS is used as reference. (c) Location where pMH94Apra-MAB_3167c plasmid integrated into the genome. Blue arrow indicates reference gene for localization. Red arrow indicates *glby* is found at the *attB* insertion site.

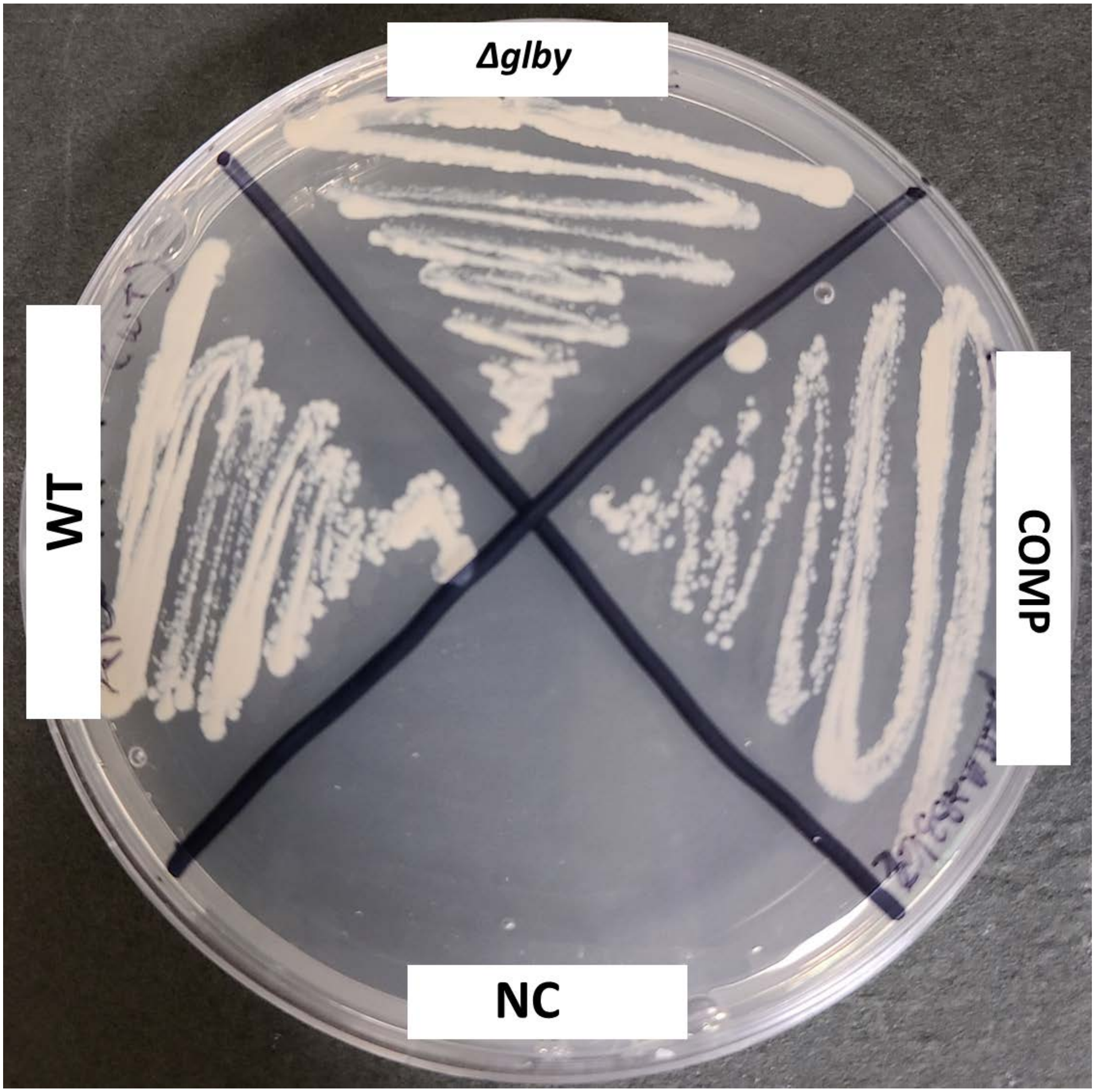


**Figure S6. Colony morphology on 7H10 agar plates**. Parent *Mab* strain ATCC 19977, Δ*glby* and complemented strain (COMP). Region of agar that was not inoculated is designated as negative control (NC).

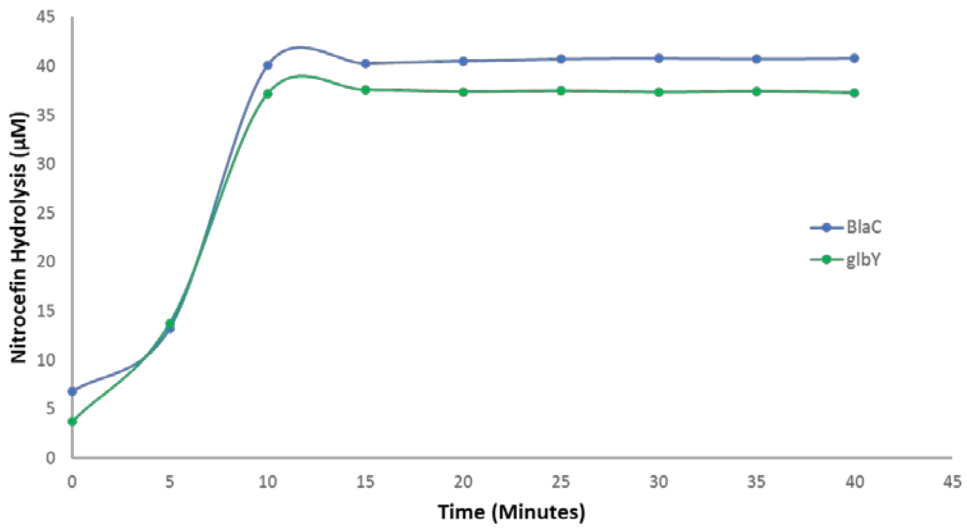


**Figure S7. Nitrocefin hydrolysis rate of Glby.** As a control comparator, a known β-lactamase, BlaC, of *M. tuberculosis* was included.

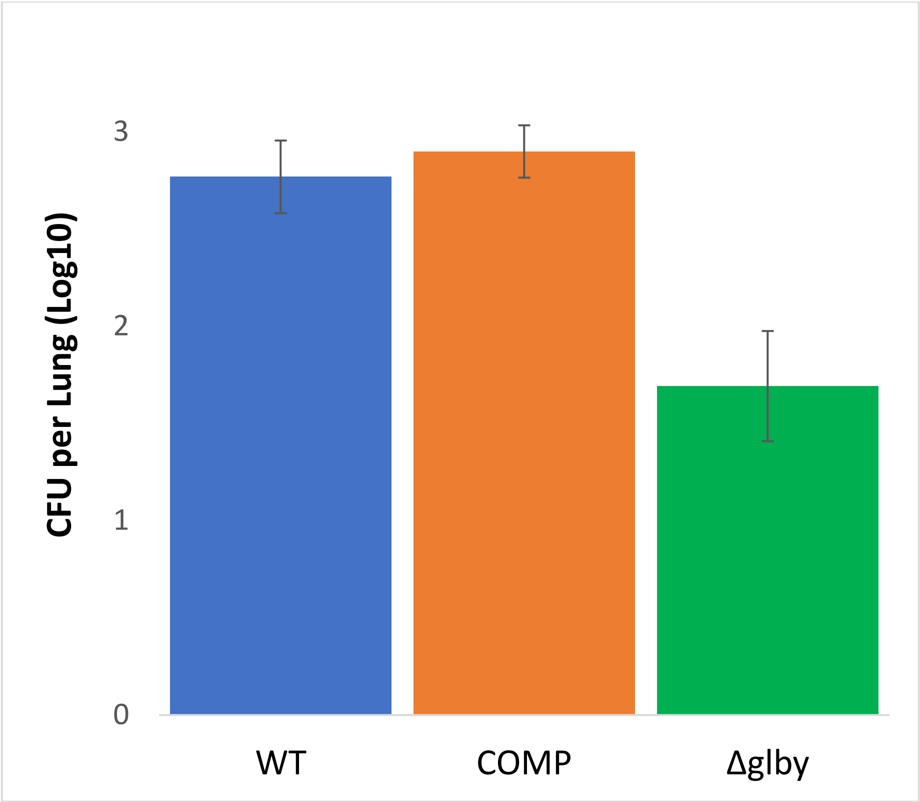


**Figure S8. Pilot Study for implantation rate of strains in the lungs of mice.** All mice were infected with aerosol of suspensions at OD ($A_{600nm}$=0.1) prepared from cultures of ATCC 19977 (WT), *Δglby* and completmented (COMP) strains at logarithmic phase of growth.
